# Stochastic competitive release and adaptive chemotherapy

**DOI:** 10.1101/2022.06.17.496594

**Authors:** J. Park, P.K. Newton

## Abstract

We develop a finite-cell model of tumor natural selection dynamics to investigate the stochastic fluctuations associated with multiple rounds of adaptive chemotherapy. The adaptive cycles are designed to avoid chemo-resistance in the tumor by managing the ecological mechanism of *competitive release* of a resistant sub-population. Our model is based on a three-component evolutionary game played among healthy (H), sensitive (S), and resistant (R) populations of *N* cells, with a chemotherapy control parameter, *C*(*t*), used to dynamically impose selection pressure on the sensitive sub-population to slow tumor growth but manage competitive release of the resistant population. The adaptive chemo-schedule is designed based on the deterministic (*N* → ∞) adjusted replicator dynamical system, then implemented using the finite-cell stochastic frequency dependent Moran process model (*N* = 10*K* – 50*K*) to ascertain the size and variations of the stochastic fluctuations associated with the adaptive schedules. We quantify the stochastic fixation probability regions of the *R* and *S* populations in the *HSR* tri-linear phase plane as a function of the control parameter *C* ∈ [0, 1], showing that the size of the *R* region increases with increasing *C*. We then implement an adaptive time-dependent schedule *C*(*t*) for the stochastic model and quantify the variances (using principal component coordinates) associated with the evolutionary cycles for multiple rounds of adaptive therapy, showing they grow according to power-law scaling. The simplified low-dimensional model provides some insights on how well multiple rounds of adaptive therapies are likely to perform over a range of tumor sizes if the goal is to maintain a sustained balance among competing sub-populations of cells so as to avoid chemo-resistance via competitive release in a stochastic environment.

## I. INTRODUCTION

The emergence of chemo-resistance in a tumor is the leading cause of progression to metastatic disease, and one of the biggest challenges facing oncologists in treating cancer patients [1–5]. The idea of managing resistance to chemotoxins in a tumor via adaptive scheduling [6, 7] emerged relatively recently from lessons learned in the field of evolutionary ecology where the goal of pest eradication has been replaced with the idea of pest management using multi-pesticide adaptive dosing schedules that are actively monitored, managed, and regulated [8, 9] so as to sustain ongoing competition among a multitude of insect species to avoid selecting for a resistant sub-type [10]. The idea that pesticides impose selection pressure on the sensitive pest population, thereby selecting for a resistant strain is by now well established [8]. The mechanism by which this process takes hold is termed competitive release [11, 12] and stems from classic observations of Connell [13] in 1961 of two species of barnacles competing for space in the intertidal zone off the Scottish coast. In this study, he noted that the blue barnacles, Balanus, occupied the intertidal zone, while the brown barnacles, Chthamalus, occupied the coast above high tide. Contrary to the commonly held belief that each occupied their own niche because of different adaptations to local micro-conditions, Connell hypothesized that it was not micro-environmental adaptation, but inter-species competition that was the dominant mechanism of selection in that region. By removing the blue barnacles from the intertidal zone, he then documented the subsequent penetration of Chthamalus into the region, concluding that the reason they now flourished was release from competition, as opposed to micro-local adaptation. A natural follow-up to this observation in situations where the goal is to reduce the dominant population (Balanus) without allowing the suppressed population (Chthamalus) to take hold was to develop sophisticated techniques to avoid competitive release and dynamically manage the competition between the competing species. The more active the management becomes, the more accurate it is to characterize the ambitious enterprise as an ongoing attempt to steer evolution [14, 15]. But to sustain this successfully over time, it has been noted that high-fidelity monitoring of the subpopulations is one of two necessary key ingredients [16], the other being the ability to accurately suppress specific sub-populations with targeted interventions. In general, since the process of steering the evolution involving multiple groups of competing sub-populations is highly stochastic [17], one could argue that the effort should be aimed at monitoring and controlling the multitude of stochastic fluctuations that result from these complex interactions [16–18]. Designing adaptive techniques to maintain these interactions becomes an attractive option, and incorporating lessons and techniques from the field of adaptive control theory, which relies heavily on the ability to collect information from sensors, then process and send the information to actuators, seems an obvious, yet non-trivial step given the many known challenges associated with controlling high-dimensional stochastic dynamical systems [19] in general, particularly in the context of evolution via natural selection in a tumor [17, 18] where sensing (sub-population monitoring) and actuating (targeted therapies) is a challenge.

Nonetheless, in the context of chemotherapy design, a multitude of targeted chemo-toxic drugs have been developed, and resistance mechanisms are becoming increasingly well understood [20, 21]. For our purposes, we are interested in approaches to manage resistance via adaptive and combination drug-scheduling, which has been pioneered by Gatenby [6, 22, 23] in oncology, and others in the related fields of controlling antibiotic evolution in microbial populations [14, 24, 25]. The complications associated with multi-drug combinations [24], drug-cycling and tailored drug-scheduling [26] has motivated us and others to undertake an effort at modeling aspects of these complex processes by developing simplified lower-dimensional mathematical/computational models in which new techniques can be tested in targeted ways, and clear lessons can be drawn from these simplied systems.

In this spirit, we introduce a model based on a finite-cell three-component stochastic evolutionary game that has well-understood [27, 28] connections to the deterministic adjusted replicator dynamical system in the infinite cell (*N* → ∞) limit. The goal, and challenges, of what we are attempting to accomplish are depicted in the schematic shown in figure 1. On the one-hand, we first design an adaptive chemotherapy schedule for the deterministic infinite-dimensional replicator system that produces a closed evolutionary cycle of sustained competition between a chemo-sensitive subgroup (S), and a chemo-resistant subgroup (R). Because the cycle is closed (i.e. periodic), it can be sustained indefinitely by repetition. This is represented in the outer loop *ABCA* shown in figure 1. In reality, however, a tumor is made up of a finite number of cells competing over an evolving fitness landscape, hence their dynamics are stochastic, and on any single realization, unpredictable fluctuations are inevitable. The discrete grid (inner triangle) associated with the Healthy-Sensitive-Resistant finite-cell phase space is also depicted in figure 1 where both the closed deterministic loop and a stochastic random walk realization are shown. Our goal in this paper is to quantify the fluctuations associated with these adaptive schedules as a function of cell number *N*, in order to quantify the uncertainty associated with the adaptive chemotherapy schedules.

**FIG. 1.**
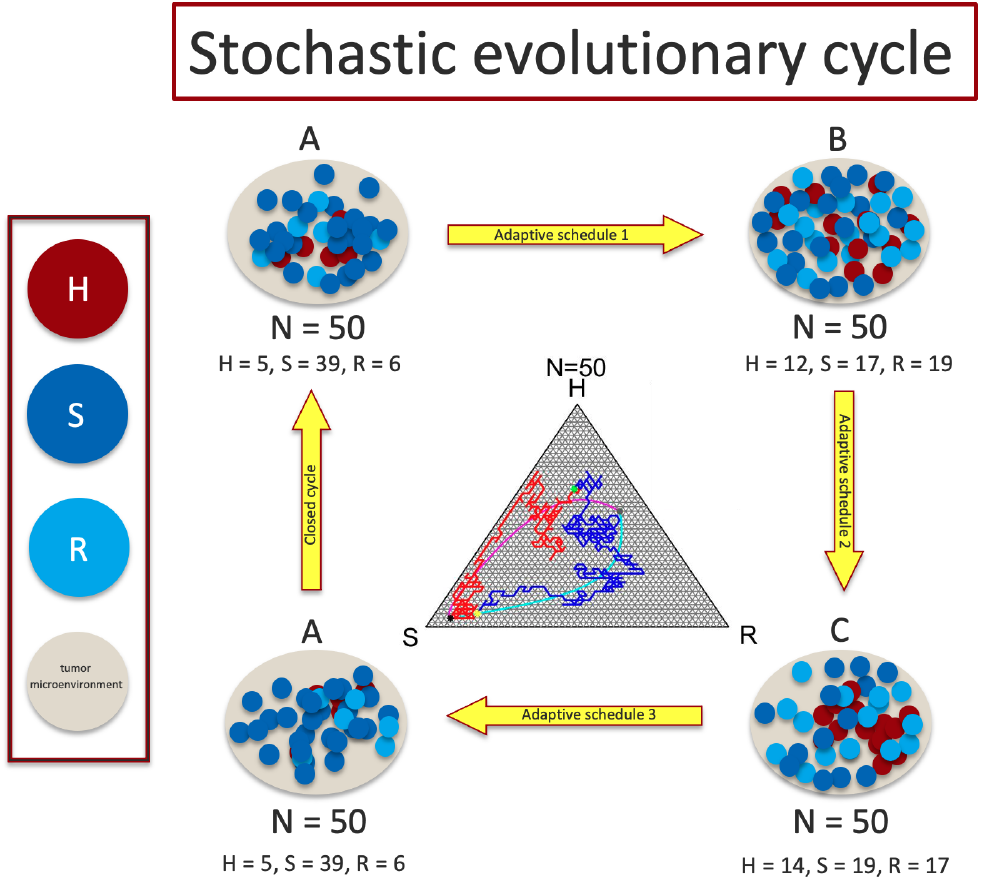
Schematic diagram describing the deterministic adaptive schedule applied to the stochastic finite-cell three-component (*H, S, R*) model. The adaptive schedule is designed to manage tumor resistance and is based on the *N* → ∞ deterministic model. Shown in the center is a discrete lattice based on the *N* = 50 cell model, with a stochastic realization attempting to mimick a closed evolutionary cycle through one round of adaptive chemotherapy.

## II. MODEL

### A. Model elements

We start by laying out the main elements of the dynamical model, depicted schematically in figure 1:

1. First we implement a three-component adjusted replicator model to track the frequencies of three sub-populations of cells: Healthy (H), Sensitive (S), and Resistant (R) [29], where *H* + *S* + *R* = 1. The cells compete in a well-mixed setting, with a payoff matrix that assigns a higher fitness to the sensitive cell population (in the absence of chemotherapy), and in addition assigns a cost to the resistant cell population [30, 31]. In the absence of chemotherapy, the sensitive population will outcompete the resistant population and saturate the tumor.
2. We use a time-dependent control function *C*(*t*) ∈ [0, 1] to model/implement the chemotherapy dosing schedule [29, 32, 33]. The function is embedded into the selection pressure parameter *w^S^*(*t*) that acts on the *S* sub-population, altering its relative fitness levels. When *C* > 0, the selection pressure on the sensitive population is lowered, allowing the resistant population to outcompete the sensitive population (competitive release) [34] if it exceeds a certain threshold. By turning the chemotherapy parameter off (*C* = 0) and on (*C* > 0), we can control the relative fitnesses of the sensitive and resistant populations in such a way to maintain their balance, without either populations saturating the tumor. By doing this adaptively, while monitoring the population balance [16], we can design schedules (0 ≤ *C*(*t*) ≤ 1) that produce closed loops in the *HSR* phase space, which we call evolutionary cycles [29]. Once the first adaptive schedule is designed, it can be indefinitely repeated to maintain the balance of the competing sub-populations.
3. We then use these adaptive schedules on a stochastic finite-cell (*N* < ∞) frequency dependent Moran process model, which in the limit N → ∞ converges to the deterministic adjusted replicator system from which the closed evolutionary cycle was designed [27, 28]. Because the model is a stochastic Markov process, the deterministic chemotherapy schedule will not necessarily reproduce the closed cycle in the finite-cell setting due to stochastic fluctuations. As a result, we seek to quantify the uncertainty associated with multiple rounds of adaptive chemotherapy [35, 36].

More details of the model are described next, with background material described in Park [33] where the model was developed and explored.

### B. Three sub-population adjusted replicator dynamics with control

We first describe the response of the *H, S*, and *R* subpopulations to the control function *C*(*t*) in the deterministic adjusted replicator model. The three sub-population frequencies are denoted:

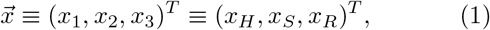

where *x_H_* + *x_S_* + *x_R_* = 1. The adjusted replicator dynamical system governing the interaction of the three groups is given by:

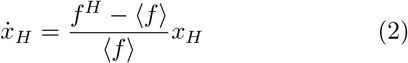

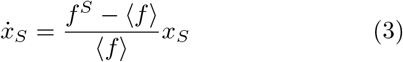

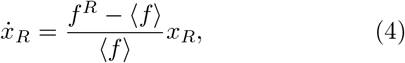

where the fitness functions of each is:

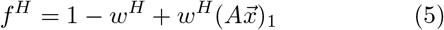

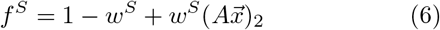

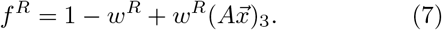

The average fitness of the entire population is then:

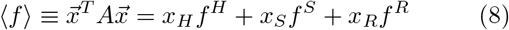

Here *A* is the 3 × 3 payoff matrix that defines the interactions of the subgroups:

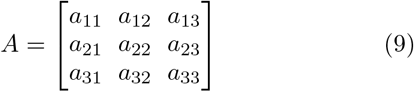

The selection pressure parameters are given by:

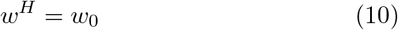

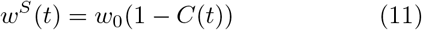

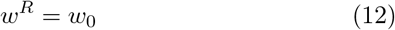

and we choose *w*_0_ = 1 (which only serves to scale time). We impose the following restrictions on the entries of *A*:

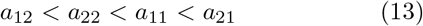

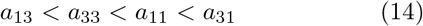

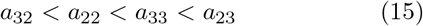

This ensures:

i. In the absence of chemotherapy (*C* = 0), we enforce a fitness cost to resistance [25], which means the sensitive population will outcompete the resistant population.
ii. In the absence of chemotherapy, the sensitive population will saturate the tumor.
iii. With chemotherapy turned on (*C* > 0), because of (11), the selection pressure on the sensitive population is reduced, allowing the resistant population to outcompete the sensitive population (if above a critical threshold).

For definiteness, we choose the specific payoff matrix values:

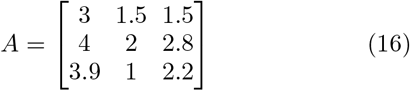

in our simulations. More details of the replicator dynamical system chemotherapy model (i.e. without normalizing the right hand sides of (2) - (4) by the average fitness term (8) can be found in [29].

### C. Three sub-population frequency-dependent Moran process with control

For a finite number of cells (*N* < ∞), we can formulate the same model as a discrete-time finite-state Markov (birth-death) process [37] that takes place on a grid in a tri-linear phase space as shown in figure 2 for *N* = 10, 20, 30, 40. One stochastic realization of the random walk that takes place on a *N* = 50 grid is depicted in figure 1. The transition probabilities for this stochastic process are given by:

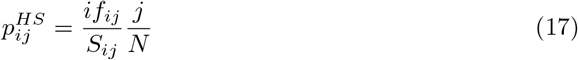

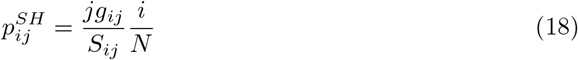

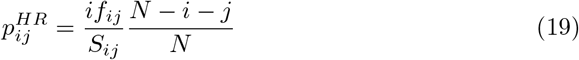

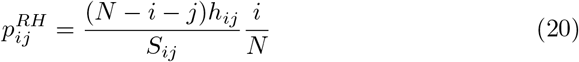

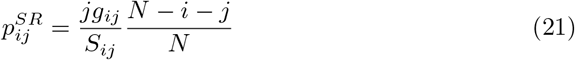

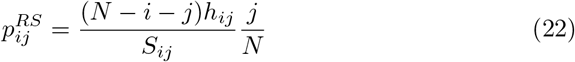

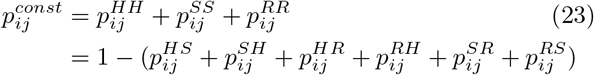

**FIG. 2.**
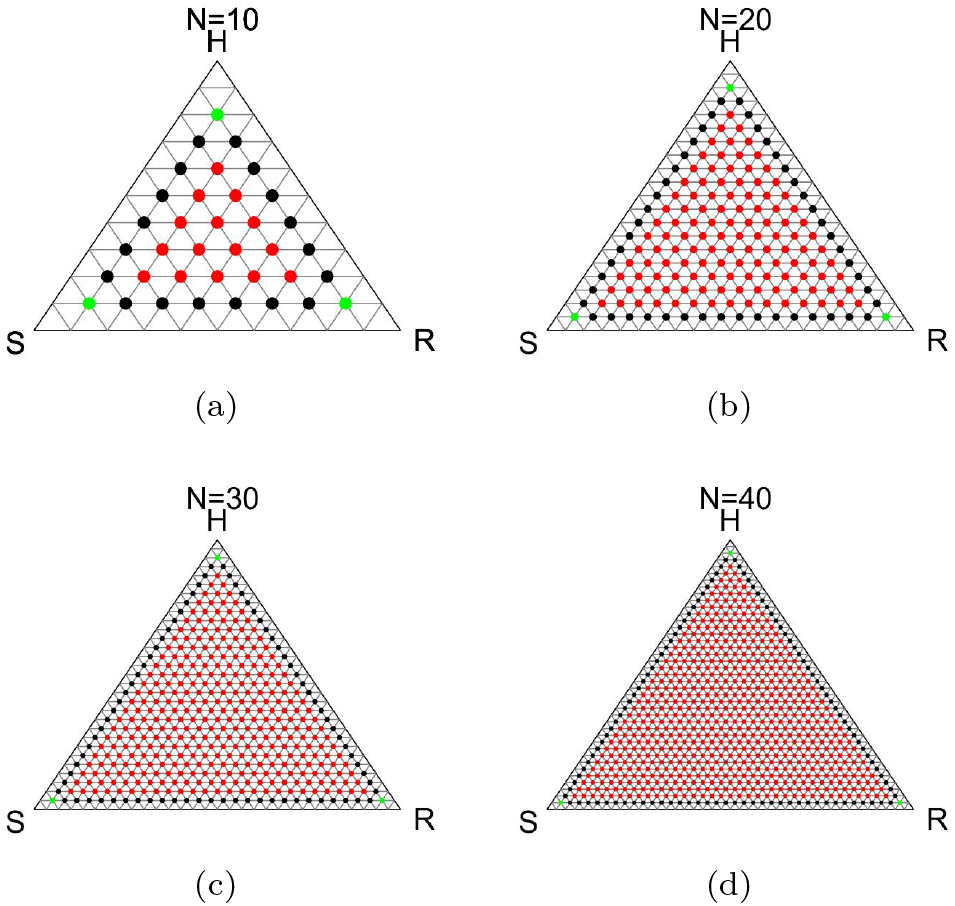
Panel showing discrete lattice associated with the finite-cell Moran process model (*N* = 10, 20, 30, 40) where each state, (*H, S, R*) = (*i, N – i – j, j*), is assigned to a lattice point. The index denoted by *i* goes along the *SH* side (*i* = 0 corresponds to the *S* corner), and the index denoted by *j* goes along the SR side (*j* = 0 corresponds to the *S* corner).

These are obtained as weighted frequencies associated with the sub-populations, weighted by the fitness functions:

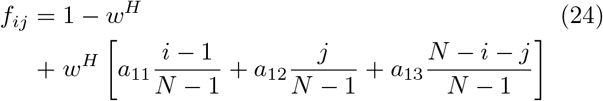

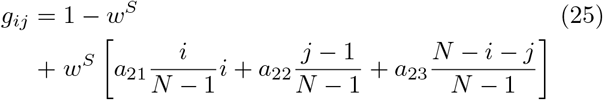

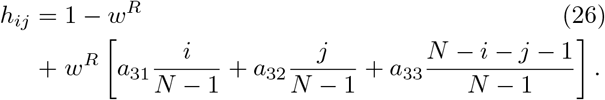

The index *i* denotes the number of cells of type *H, j* denotes the number of cells of type *R*, and *N* – *i* – *j* denotes the number of cells of type *S*, with 0 ≤ *i* ≤ *N*; 0 ≤ *j* ≤ *N, i* + *j* ≤ *N*. The state of the system is given by (*i, j*) (visualized as a grid point on a triangular grid as in Figure 2) with probability of residing in that state after *n* steps given by 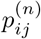.

As in the continuous model, the parameter *w^α^* ∈ [0, 1] (*α* = *H, S, R*) controls the selection pressure and allows us to independently adjust the fitness landscape. At the two extremes, when *w^α^* = 0, the payoff matrix makes no contribution to fitness (only random drift [37] governs the dynamics). At the other extreme, when *w^α^* = 1, the payoff matrix makes a large contribution to the fitness, with selection pressure driving the dynamics. We define the average tumor fitness, 〈*f*〉_*N*_, as the discrete analogue of (8):

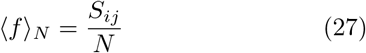

where:

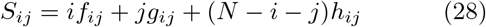

To implement chemotherapy in the discrete Markov process, we discretize time 0 ≤ *t* ≤ *T* in (2)–(4) so that at each time-step n in the process (with *n* = *tN*), we enforce the time-dependent chemotherapy schedules by adjusting the selection pressures, where:

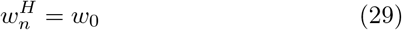

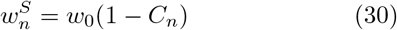

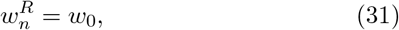

This renders the Markov process described in [38, 39] non-stationary [40].

### D. The mean-field limit of infinite populations

To make the connection between the finite-cell (*N* < ∞) Moran process model and the deterministic adjusted replicator dynamics equations (*N* → ∞), we limit ourselves in this section to a description of the two subpopulation system to minimize notational complexity, but we note that the three and higher sub-population case can be treated in an identical way, albeit with more notational overhead. We follow the derivation layed out in [27, 28] in condensed form.

For the two sub-populations, *A*, and *B*, let *i* be the number of individuals in sub-population *A*, let *j* be the number of individuals in sub-population *B* with *i* + *j* = *N*, and let 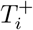 be the transition probability from state *i* to *i* + 1. Let 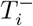 denote the transition probability from state *i* to *i* – 1. Then 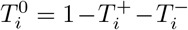 denotes the probability that the system stays at its current state (under the assumption of no mutations). In this model, mutations have already accrued before chemotherapy cycles begin, resulting in the resistant sub-population. Both *i* = 0 and *i* = *N* are absorbing states corresponding to a homogeneous B population and a homogeneous *A* population, respectively, and these lead to 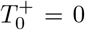 and 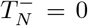, or 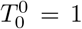 and 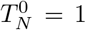. For all other states, the transition probability, 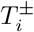 is computed by considering the birth-death process. The birth rate is assumed to be proportional to the expected fitness, 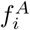 and 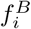, of *A* and *B* which are, in turn, functions of the expected payoffs, 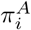 and 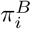:

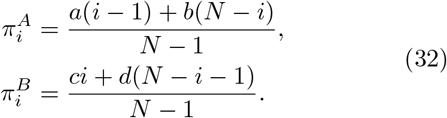

To compute these payoffs, we use the 2 × 2 payoff matrix:

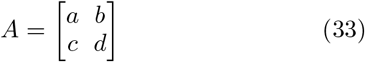

in the context of an evolutionary game [37]. The fitness functions are defined by:

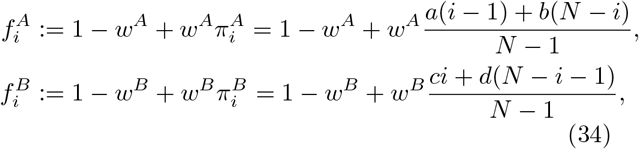

where 0 ≤ *w^A^* ≤ 1 and 0 ≤ *w^B^* ≤ 1 are selection intensity parameters. *w_A_* ≪ 1, *w_B_* ≪ 1 are the weak selection limits [37] of A and B, respectively. To compute the transition probabilities 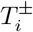, we use fitness-weighted frequency-dependent averages 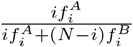 and 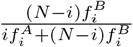 as birth rates of the *A* and *B* sub-populations respectively, which then give rise to transition probabilities:

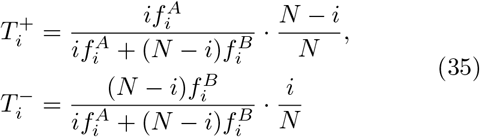

for *i* ∈ {1, ···, *N* – 1}. Taken together, we can now write the (*N* + 1) × (*N* + 1) tridiagonal transition matrix T governing the Markov process:

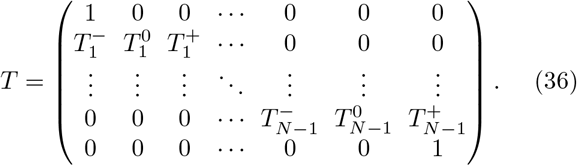

We can then formulate the process in terms of the probability, *P_i_*(*τ*), that the system is in the state *i* at time *τ*, with the transition probabilities:

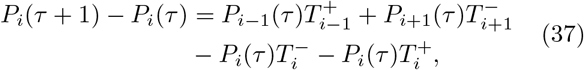

which is called the *the master equation* [41]. It describes the net change in probability from state *i* in one time step from time, *τ*. Letting

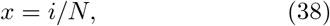

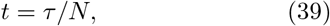

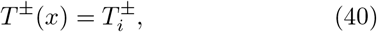

it can be rewritten in terms of the probability density function, *ρ*(*x,t*) ≡ *NP_i_*(*τ*):

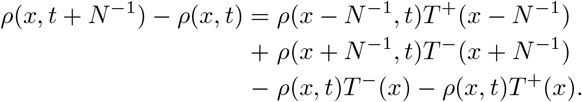

For large *N*, the Taylor expansion at *x* and *t* for both the probability density function, *ρ*, and the transition probabilities, *T*^±^, up to the first order term in *N*^−1^, gives rise to *the Kolmogorov forward equation* [41]:

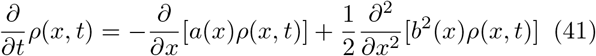

with a drift term, *a*(*x*) = *T*^+^(*x*) – *T*(*x*), and a diffusion term, 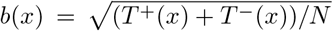. Equation (41) is equivalent to the stochastic differential equation (via Itô calculus):

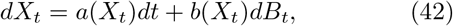

which is the governing *the Langevin equation* [41]. Here *B_t_* is the one dimensional standard *Wiener process* and *X_t_* is the state of the system at time *t*. Noting that the diffusion term, *b*(*X_t_*), vanishes with the rate of 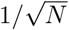 as *N* → ∞, the limiting system is solely determined by the drift term, *a*(*X_t_*), as follow:

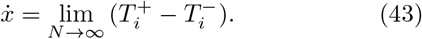

From equations in (35), this becomes

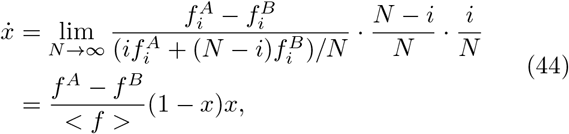

i.e. the adjusted replicator equation recovered, in the limit of *N* → ∞ from the master equation, for the Moran process with *x* = *x_A_*. More detailed derivations are available in [27, 28] even when the Moran process is applied to a population of *N*, sub-divided into *d* > 2 subpopulations. For our purposes, the brief description is sufficient to establish the connection between the finite-cell (*N* < ∞) frequency-dependent Moran birth-death process and the deterministic adjusted replicator dynamical system in passing to the limit *N* → ∞.

## III. RESULTS

### A. Stochastic competitive release

First we detail the process of competitive release of the resistant sub-population (R) as a function of the chemotherapy parameter *C* ∈ [0, 1]. It is straightforward to prove that for *C* < 1/3, all trajectories converge to the *S* corner, and if *C* > 1/2, all trajectories converge to the *R* corner. This is shown in figure 3(a) and (d). For values 1/3 ≤ *C* ≤ 1/2, the basins of attraction of the two corners are intermingled as shown in figure 3(c) for *C* = 0.45. The fixed point is located at:

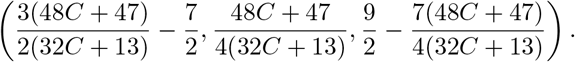

**FIG. 3.**
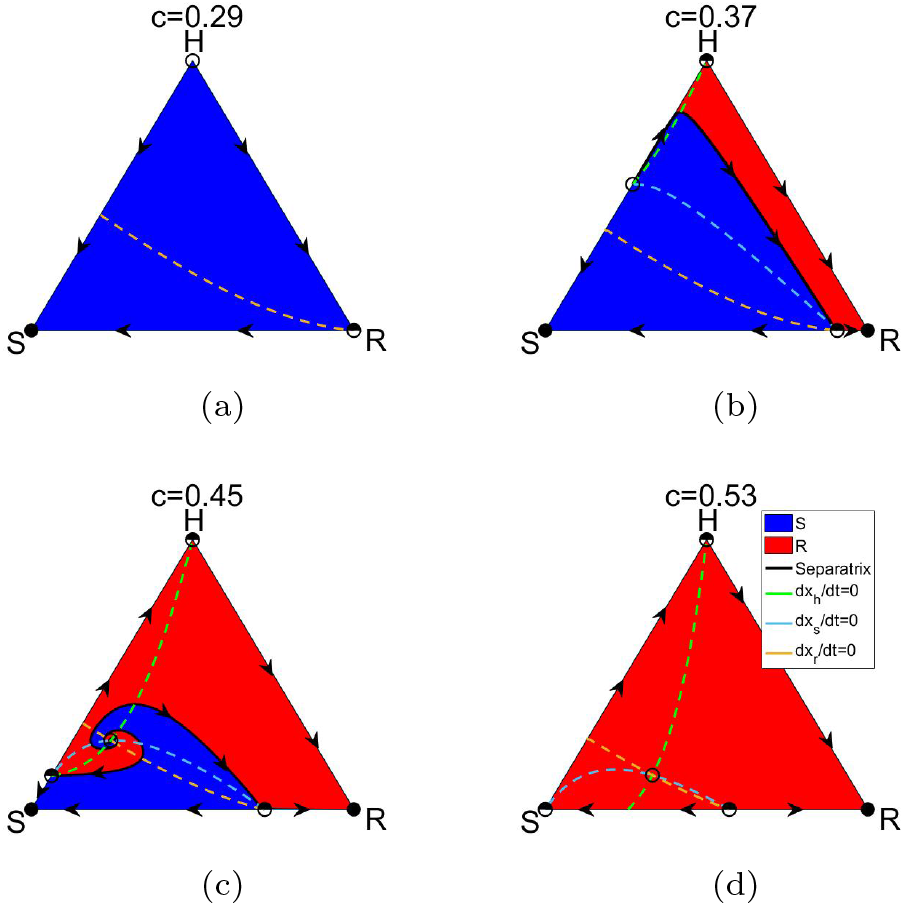
Basins of attraction and nullclines associated with the deterministic adjusted replicator system with three subpopulations (*H, S, R*). Panel shows the separatrices and nullclines (*dx_H_/dt* = 0, *dx_S_/dt* = 0, *dx_R_/dt* = 0) through the balanced fitness interior fixed point that determines the basin boundaries of attraction associated with the *S* state and the *R* state. The interior fixed point represents the state at which each of the subpopulation fitness levels exactly matches the average fitness of the entire population. Filled circles are stable, unfilled circles are unstable. (a) *C* ≡ 0.29; (b) *C* ≡ 0.37; (c) *C* ≡ 0.45; (d) *C* ≡ 0.53

For values 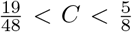, the basins of attraction associated with the *S* and *R* corners spiral out from the internal fixed point, as shown in figure 3(c). The process of competitive release unfolds if the initial conditions associated with the competing sub-populations are in the red region. For values of the chemotherapy parameter large enough (*C* = 0.53 shown in figure 3(d)), all trajectories converge to the R corner – the resistant population is the evolutionarily stable state (ESS) [37]. Figure 3 also depicts the nullclines (*dx_H_*/*dt* = 0, *dx_S_/dt* = 0, *dx_R_/dt* = 0) in the three-component phase-space which divides the triangluar region into further dynamical subsections. Exactly how the stochastic version of competitive release tracks the deterministic one is depicted in the panel shown if figure 4 (for *N* = 100, 500, 1000, 2000). For clarity, we overlay the sharp deterministic separatrices on top of the stochastic boundaries. Colors depict the probability of convergence to the R corner, with the color panel shown on the right. Red indicates convergence to R with probability 1, blue indicates convergence to R with probability 0 (equivalent to convergence to S with probability 1). Surrounding the deterministic separatrices is the stochastic transition layer with probabilities between 1 and 0. Details of the shape of the boundary emerge with larger values of N, particularly near the central spiral associated with the internal unstable fixed point where the subpopulation fitness levels equal the average fitness of the full system. The sequence shown in figures 4(c),(g),(k), and (o) most clearly show the stochastic spiral emerging with increasing *N* for *C* = 0.45.

**FIG. 4.**
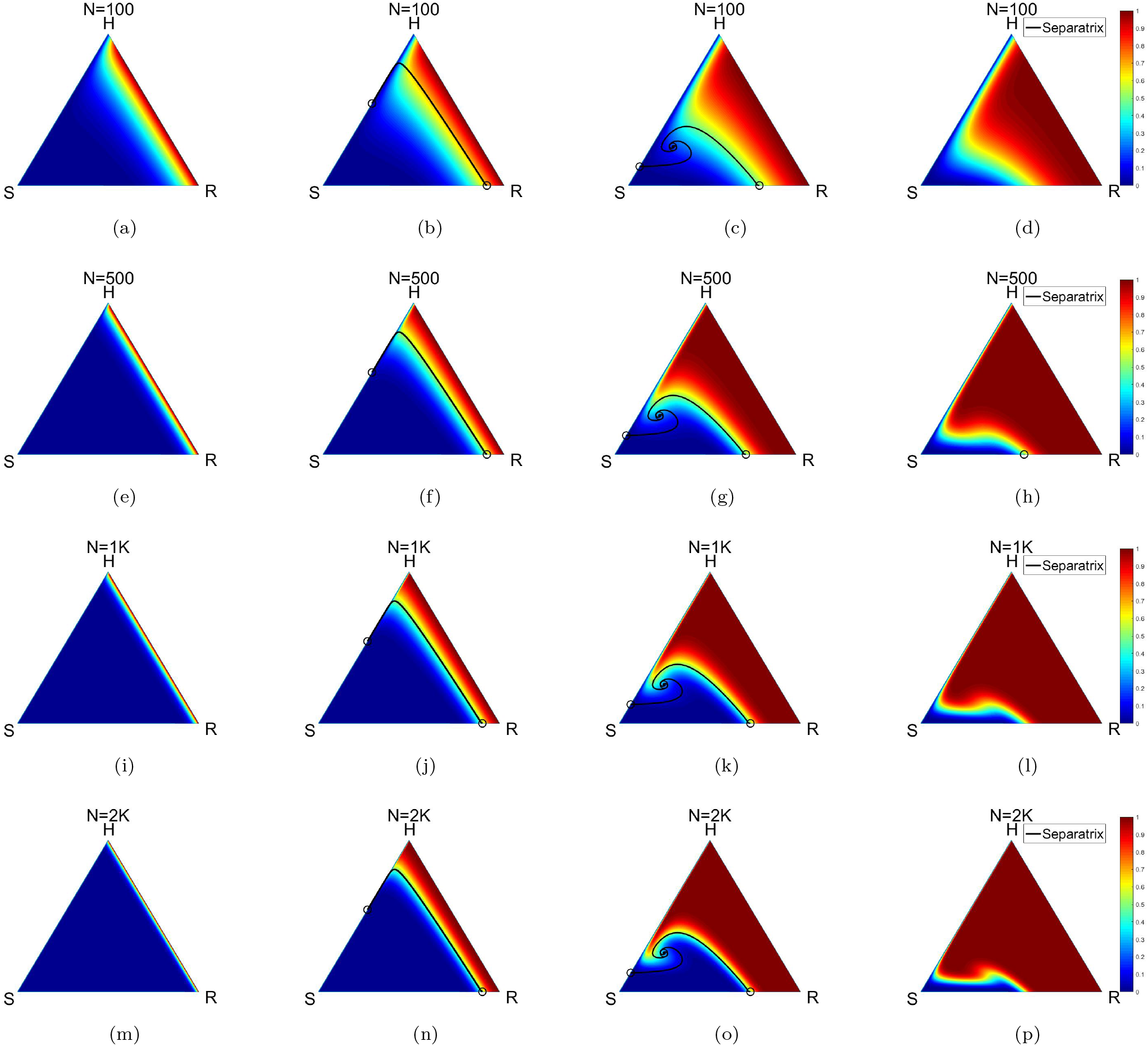
Panel shows the the fixation probability associated with being absorbed in the R state (red corresponds to probability 1, blue corresponds to probability 0) with the (sharp) deterministic (adjusted replicator dynamics) separatrix overlaid on the diagrams. As *N* increases, the stochastic boundary sharpens and coincides with the deterministic boundary. (a) *C* ≡ 0.29; (b) *C* ≡ 0.37; (c) *C* ≡ 0.45; (d) *C* ≡ 0.53; (e) *C* ≡ 0.29; (f) *C* ≡ 0.37; (g) *C* ≡ 0.45; (h) *C* ≡ 0.53; (i) *C* ≡ 0.29; (j) *C* ≡ 0.37; (k) *C* ≡ 0.45; (l) *C* ≡ 0.53; (m) *C* ≡ 0.29; (n) *C* ≡ 0.37; (o) *C* ≡ 0.45; (p) *C* ≡ 0.53.

### B. Deterministic adaptive cycles

We turn next to adaptive chemotherapy using time-dependent chemotherapy parameter *C*(*t*). Shown in figure 5 are deterministic trajectories associated with two values of the chemotherapy parameter. Figure 5(a) shows convergence to the evolutionarily stable *S* corner with *C* = 0, while figure 5(b) shows convergence to the *R* corner with *C* = 0.7. The key idea behind constructing adpative schedules, as described thoroughly in [29], is to overlay these two families of trajectories, as shown in figure 5(c). What becomes clear with these overlaid trajectories is there are many families of closed loops that can be created by switching between *C* = 0 and *C* = 0.7 at the times the orbits cross. One of these closed loops, which we label *PQ*, is shown in figure 5(d). Once this *PQ* loop is identified, and the crossing times are calculated, the loop can be repeated indefinitely by repeating the chemotherapy schedule of switching from *C* = 0.7 (traversing from *P* to *Q*), to *C* = 0 (traversing back from *Q* to *P*). The process and many examples are discussed in much more detail in [29]. We are interested in properties associated with stochastic versions of these cycles produced by overlying the stochastic versions of these two families of trajectories, as shown in the panel in figure 6. For any finite value of *N*, the deterministic and stochastic trajectories are different, but as *N* gets larger, the more they track each other. Overlaying two of these stochastic trajectories, while close (in some sense) to the deterministic *PQ* loop discussed previously, will not coincide exactly. We characterize the differences next.

**FIG. 5.**
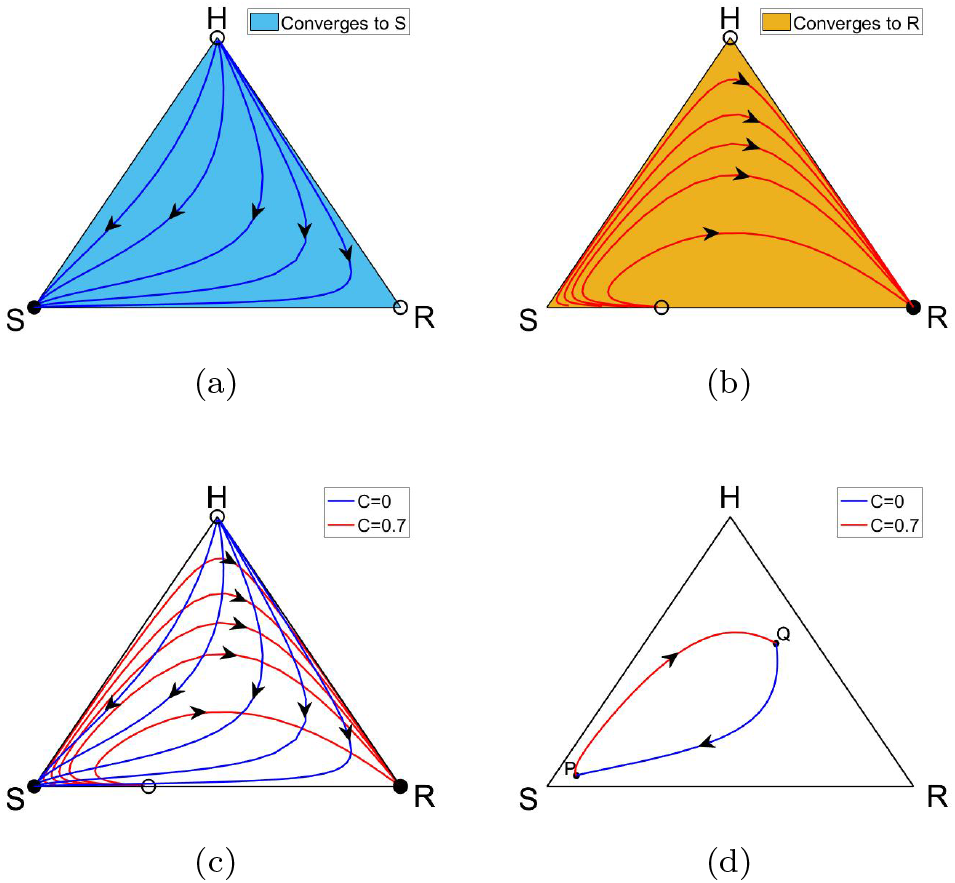
Deterministic trajectories of the adjusted replicator system for different constant chemotherapy values. (a) For *C* ≡ 0, all trajectories converge to the *S* corner; (b) For *C* ≡ 0.7, all trajectories converge to the *R* corner (competitive release); (c) If both families of trajectories are overlaid, it is clear that closed loops can be obtained if the two different values of *C* are switched from one to the other at the times at which the trajectories cross. We call these closed loops produced by time-dependent *C*(*t*) deterministic evolutionary cycles; (d) Example of a deterministic evolutionary cycle PQP obtained by setting *C* = 0.7 on the first half-cycle from P to Q, then switching to C = 0 on the second half-cycle from Q to P. The closed cycles can be repeated over and over indefinitely through multiple rounds *PQPQPQPQP*….

**FIG. 6.**
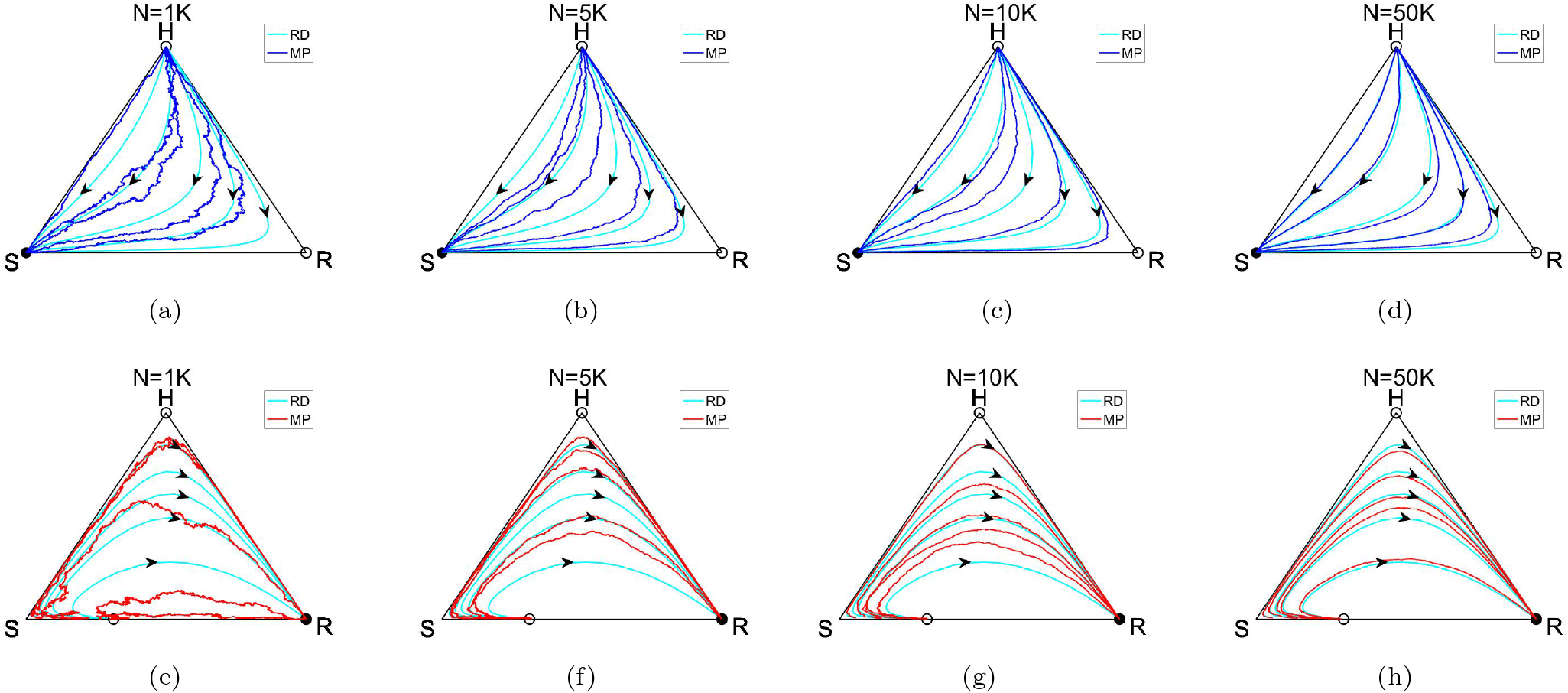
Panel depicting stochastic realizations of the finite-cell Moran process (MP) model (*N* = 1*K*, 5*K*, 10*K*, 50*K*) compared to the deterministic adjusted replicator dynamical system (RD) for *C* = 0 (a,b,c,d), and *C* = 0.7 (e,f,g,h). As *N* increases, the stochastic trajectories converge (pointwise) to the deterministic trajectories.

### C. Stochastic adaptive cycles

Figure 7 shows the spread of the distribution of points on each half-cycle (*P* to *Q*, then *Q* to *P*), for 1, 000 stochastic realizations, for *N* = 1*k*, 5*k*, 10*k*, 50*k*. Clearly the spread around the points *P* and *Q* tightens as *N* increases. This is seen more clearly in figure 8 where we plot the distributions around *P* and *Q* in both the *SR* plane, and the principal component axis planes. We employ a kernal density estimator around the points for visual clarity. It is clear from figures 8(e)-(h) that as *N* increases, the spread around *Q* (first half-cycle) can be thought of as a multivariate Gaussian distribution with means and standard deviations shown in the captions. Likewise, we show in figures 8(m)-(p) the stochastic spread around *P* (second half-cycle). Again, the distributions closely resemble a multivariate Gaussian distribution around each point.

**FIG. 7.**
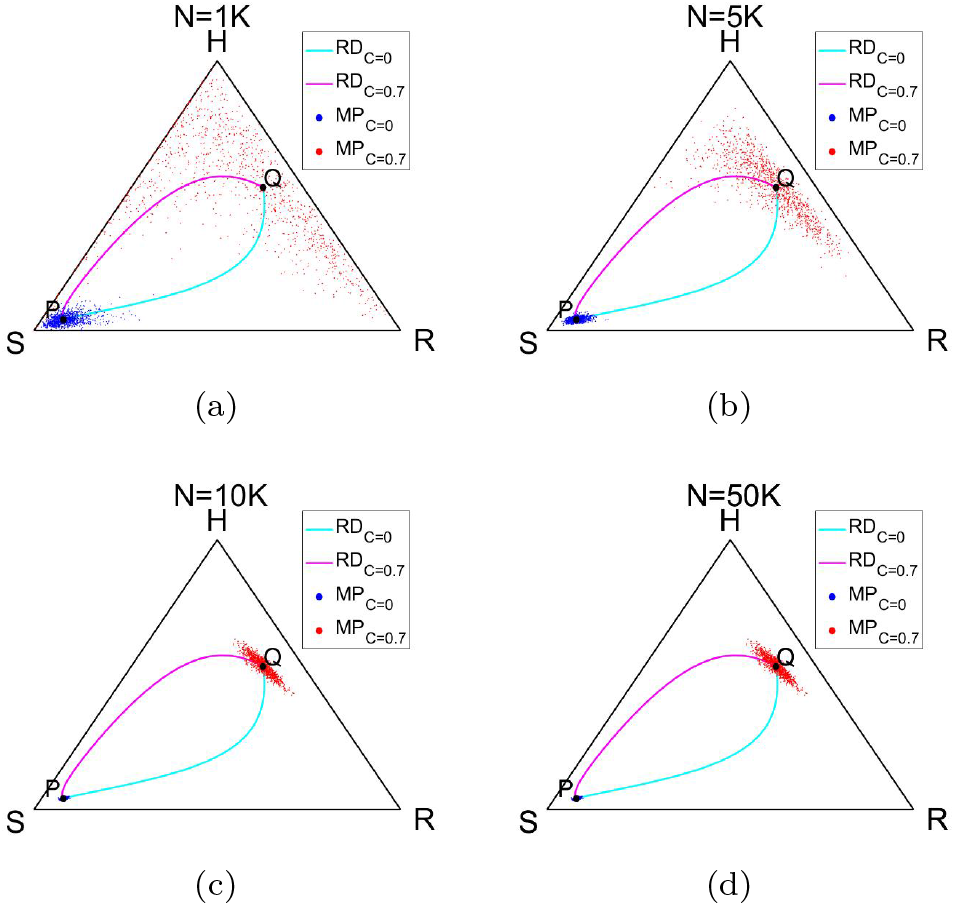
The spread of the distribution of points around *P* and *Q* for 1, 000 realizations of one stochastic adaptive cycle. The first half-cycle *PQ* uses *C* = 0.7, while the second half-cycle *QP* uses *C* = 0. (a) *N* = 1*K*; (b) *N* = 5*K*; (c) *N* = 10*K*; (d) *N* = 50*K*

**FIG. 8.**
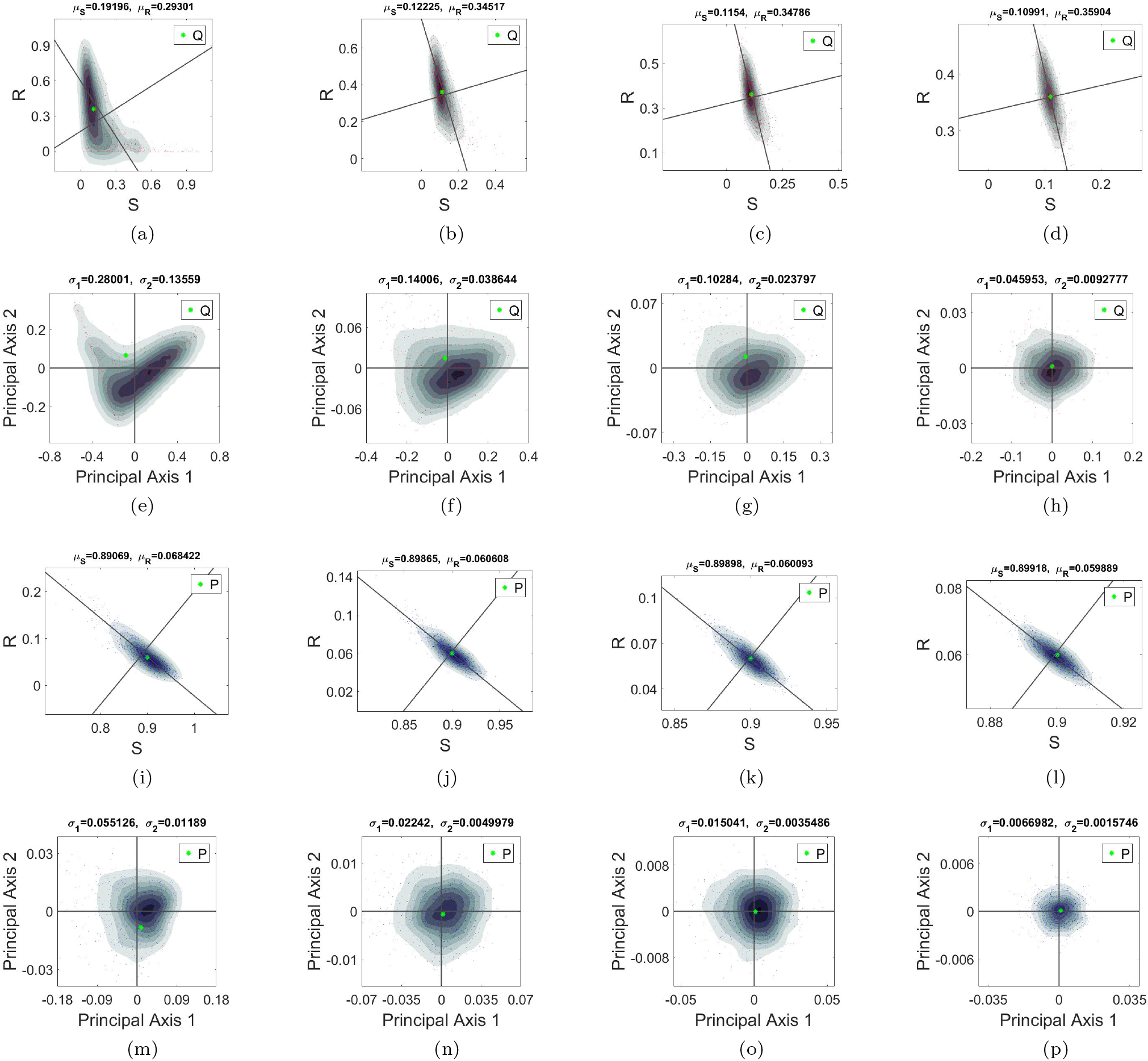
Stochastic spread (1, 000 realizations) of one evolutionary cycle *PQP* in the *SR* plane, and the principal axis plane *P*_1_*P*_2_. (a)-(h) shows the spread around the point *Q* (first half-cycle) achieved with *C* = 0.7, (i)-(p) shows the spread around the point *P* (second half-cycle) achieved with *C* = 0. The spread around each point (for sufficiently large *N*) can be thought of as an approximate multivariate Gaussian distribution (highlighted with contours from a kernal density estimator) with mean *μ_i_* and semi-major/minor axes *σ_i_*. The mean frequency, *μ_S_* (or *μ_R_*), of the *S* and *R* subpopulations around *P* and *Q* converges to the deterministic proportion of *S, R* as *N* increases, while the semi-major axis, *σ*_1_, and semi-minor axis, *σ*_2_ decrease. (a) *N* = 1*K*, *μ_S_* = 0.1920, *μ_R_* = 0.2930; (b) *N* = 5*K*, *μ_S_* = 0.1223, *μ_R_* = 0.3452; (c) *N* = 10*K*, *μ_S_* = 0.1154, *μ_R_* = 0.3479; (d) *N* = 50*K, μ_S_* = 0.1099, *μ_R_* = 0.3590; (e) *N* = 1*K*, *σ*_1_ = 0.2800, *σ*_2_ = 0.1356; (f) *N* = 5*K, σ*_1_ = 0.1401, *σ*_2_ = 0.0386; (g) *N* = 10*K, σ*_1_ = 0.1028, *σ*_2_ = 0.0238; (h) *N* = 50*K, σ*_1_ = 0.0460, *σ*_2_ = 0.0093; (i) *N* = 1*K, μ_S_* = 0.8907, *μ_R_* = 0.0684; (j) *N* = 5*K, μ_S_* = 0.8987, *μ_R_* = 0.0606; (k) *N* = 10*K, μ_S_* = 0.8990, *μ_R_* = 0.0601; (l) *N* = 50*K, μ_S_* = 0.8992, *μ_R_* = 0.05989; (m) *N* = 1*K, σ*_1_ = 0.0551, *σ*_2_ = 0.0119; (n) *N* = 5*K, σ*_1_ = 0.0224, *σ*_2_ = 0.0050; (o) *N* = 10*K, σ*_1_ = 0.0150, *σ*_2_ = 0.0035; (p) *N* = 50*K, σ*_1_ = 0.0067, *σ*_2_ = 0.0016

### D. Uncertainty quantification over multiple rounds

Because of the stochastic spread over the first *PQ* cycle, we ask how repeatable the cycles are as they are repeated round-by-round. Figure 9 shows the results of repeating the cycle through 4 rounds: *PQPQPQPQ*. By the fourth round, the spread of points (around *Q*, for example in (d) and (h)) shows very little structure. Figure 10 shows the four rounds in the *SR* plane and the principal component planes. Here, we can more clearly see (figure 10(e),(f)) the multivariate Gaussian nature of the distribution for the first two rounds, but by the third and fourth rounds (figure 10(g),(h)), the structure clearly begins to break down. In figure 11 we show the size of the principal components (standard deviations) as a function of the first eight rounds of adpative therapy, on a log-log scale, with linear curve fits shown. As the rounds increase, the spread follows power-law distributions fairly cleanly. The linear curve fits for the semi-major (minor) axis of the distribution of the points around *P* (1, 000 realizations starting at *P*) follow:

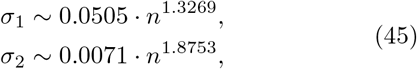

for *N* = 10*K*, and

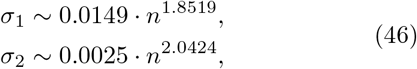

for *N* = 50*K*. Here, *σ*_1_ is the semi-major axis, *σ*_2_ is the semi-minor axis, and *n* is the number of rounds.

**FIG. 9.**
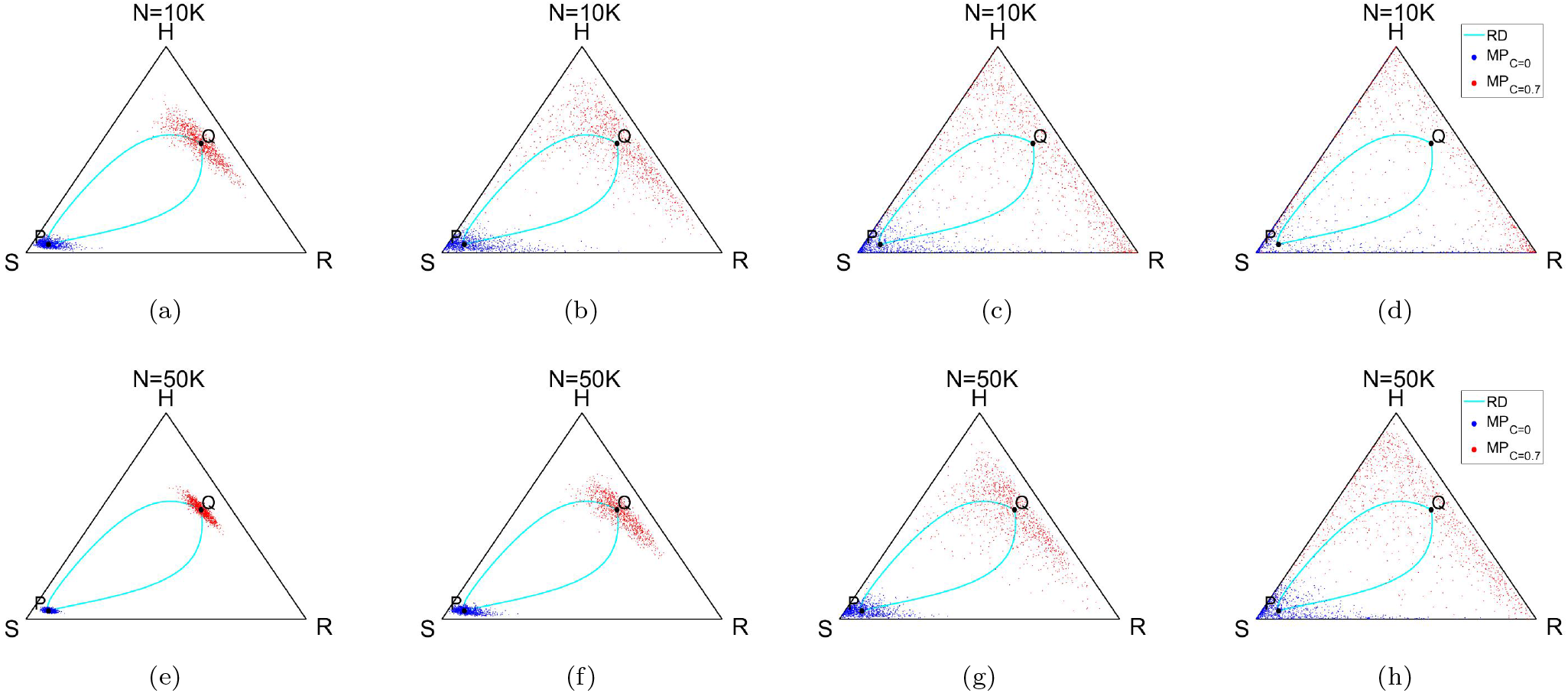
Panel depicting the spread of the stochastic evolutionary cycles *PQPQPQPQP*…, round-by-round, associated with the finite-cell Moran process model (1, 000 realizations for *N* = 10*K*, 50*K*). (a) *N* = 10*K*, round 1; (b) *N* = 10*K*, round 2; (c) *N* = 10*K*, round 3; (d) *N* = 10*K*, round 4; (e) *N* = 50*K*, round 1; (f) *N* = 50*K*, round 2; (g) *N* = 50*K*, round 3; (h) *N* = 50*K*, round 4

**FIG. 10.**
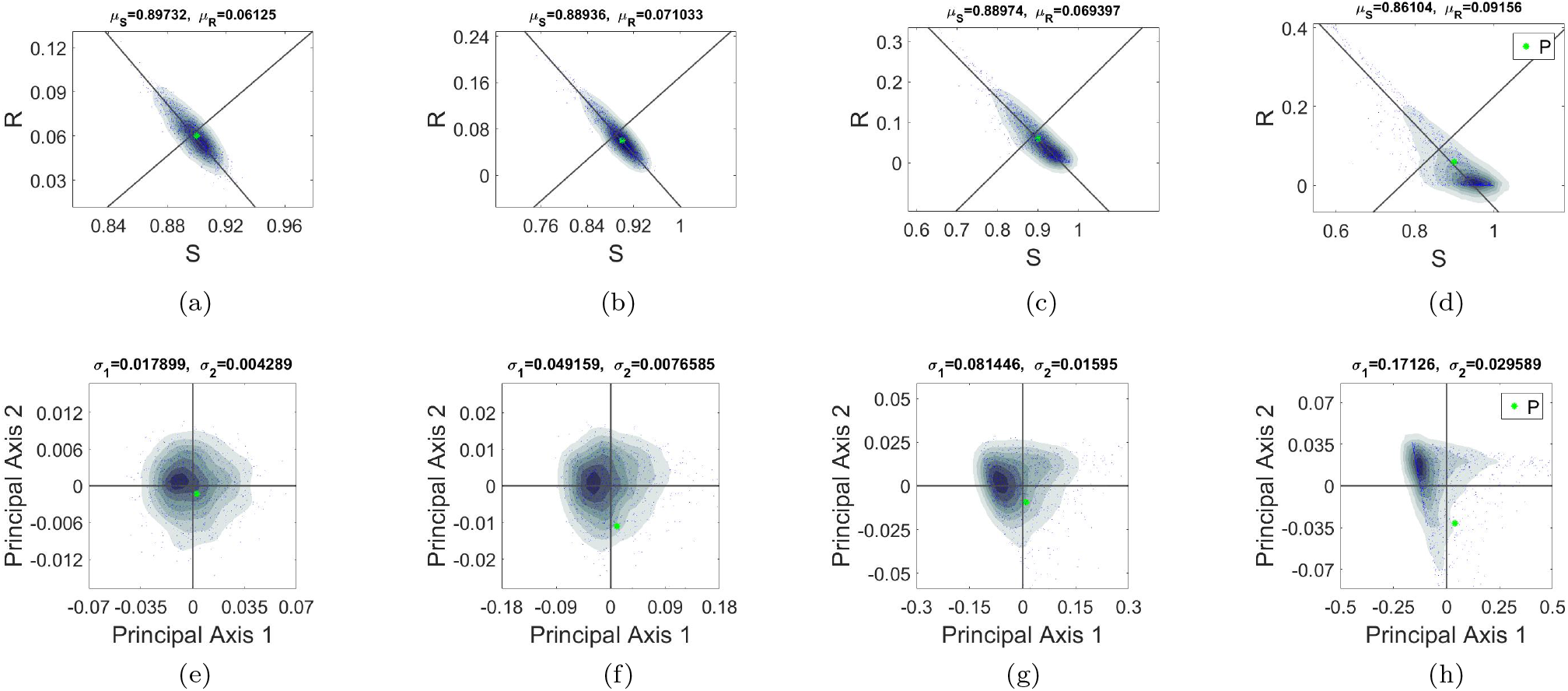
Stochastic spread (1, 000 realizations, *N* = 50*K*) around the point *P* of multiple (4 rounds) evolutionary cycles *PQPQPQPQP*… in the *SR* plane, and the Principal axis plane *P*_1_*P*_2_. As the number of rounds increases, the spread of the distribution of points around *P* becomes centered towards the homogeneous state *S* and loses the multivariate Gaussian-like symmetry. (a) Round 1, *SR* plane; (b) Round 2, *SR* plane; (c) Round 3, *SR* plane; (d) Round 4, *SR* plane; (e) Round 1, *P*_1_*P*_2_ plane; (f) Round 2, *P*_1_*P*_2_ plane; (g) Round 3, *P*_1_*P*_2_ plane; (h) Round 4, *P*_1_*P*_2_ plane

**FIG. 11.**
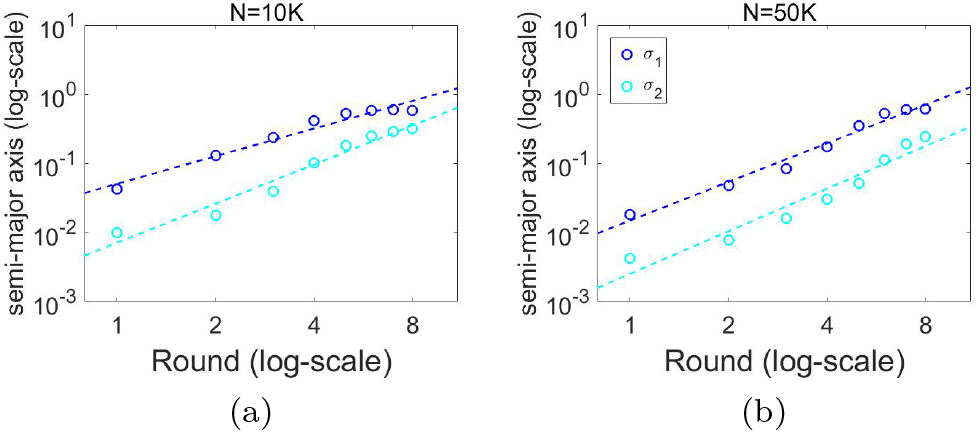
Semi-major and semi-minor axes of the distribution of points around *P* through 8 rounds of adaptive therapy cycles, 1, 000 realizations of the finite-cell Moran process with *N* = 10*K*, 50*K*, plotted on log-log scale to show power-law scaling of the accumulated stochastic error associated with applying the adaptive schedules over multiple cycles.

## IV. DISCUSSION

The goal of adaptive therapy is to leverage the competition between sensitive and resistant cells in order to delay chemo-resistance as long as possible [6]. The approach maintains a certain tumor burden in order to achieve this, hence it is sometimes criticized for its deliberate goal to maintain rather than cure. Nonetheless, there are many other diseases, such as diabetes and HIV, where a long-term maintanance approach has been transformative, and one might conjecture that a strategy of transitioning certain cancer types from an accute disease to a chronic condition that is tolerable could be successful. There are, in fact, several promising ongoing clinical trials testing adaptive concepts (NCT02415621, NCT03511196, NCT03543969, NCT03630120). For example, trial NCT02415621 longitudinaly tracks cohorts of metastatic castration resistant prostate cancer patients using two clinical biomarkers, PSA and testosterone, while using aberaterone as a therapeutic agent to attempt to manage resistance [42–44], with alterations to the treatment schedules made adaptively as the tumor changes. These trials represent a significant departure from standard chemotherapy treatment protocols such as maximum tolerated dose (MTD) schedules, and low-dose metronomic therapies [45, 46] that are pre-scheduled and take a more aggressive approach to killing every last cancer cell. But these approaches are increasingly being viewed as potentially counterproductive in many cases [47], and can actually contribute to tumor resistance and treatment failure.

Aside from clinical trials, there are certainly fundamental questions and potential modifications [48] that need to be explored associated with adaptive protocols, and some of them are best addressed using simulations. It is unclear, for example, what the optimal balance of sub-populations should be in order for adaptive methods to work best [48]. In our model, this translates to the question of what location in the triangluar phase space (shown in figure 1) is most effective for targeting. But more generally, assessing the accuracy and repeatability of adaptive chemotherapeutic dosing schedules over multiple rounds in a stochastic setting is particularly important for understanding performance over populations of patients and a range of tumor sizes and types. It is well documented that tumor heterogeneity provides fuel for resistance [49], and tumor cell heterogeneity and the effectiveness of repeated rounds of adaptive dosing schedules in a stochastic setting are presumably linked.

## V. CONCLUSION

In this paper, we describe an adaptive chemotherapy model, and develop a method of assessing the accuracy and repeatability of adaptive dosing schedules that attempt to manage the mechanism of competitive release in a stochastic setting. If one accepts the premise that evolutionary processes and selection pressure are at play in chemotherapeutic control of a tumor, then dealing with stochasticity is inevitable [17]. We do this by developing an uncertainty quantification method [35] tailored to our setting, which characterizes the size of the variation in the dosing target points through a closed evolutionary loop, over multiple rounds. We demonstrate that the accumulated stochastic error associated with applying adaptive schedules over multiple cycles follows powerlaw scaling (as opposed to exponential). Maintaining a handle on these accumulated errors caused by stochastic fluctuations over multiple adaptive chemotherapy sessions is just one of the challenges that must be overcome in order for these types of approaches to be embraced more widely.

## ACKNOWLEDGMENTS

We gratefully acknowledge support from the Army Research Office MURI Award #W911NF1910269 (2019-2024) as well as useful conversations and suggestions from Prof. Kukavica and Prof. Ziane.

